# Chronic exposure to IL6 leads to deregulation of glycolysis and fat accumulation in the zebrafish liver

**DOI:** 10.1101/2020.07.04.162008

**Authors:** Manoj K Singh, Rijith Jayarajan, Swati Varshney, Sindhuri Upadrasta, Archana Singh, Rajni Yadav, Vinod Scaria, Shantanu Sengupta, Dhanasekaran Shanmugam, Shalimar, Sridhar Sivasubbu, Sheetal Gandotra, Chetana Sachidanandan

## Abstract

**BACKGROUND AND AIMS:** Inflammation is a constant in Non-Alcoholic Fatty Liver Disease (NAFLD) and is usually considered a consequence. We propose that inflammation can be a cause for NAFLD. Obesity is strongly associated with (NAFLD), but not always. NAFLD in lean individuals is more common in certain populations, especially Asian-Indians. Lean healthy Indians also have a higher basal circulating IL6 suggesting a link with inflammation. We propose that inflammation-induced fatty liver could be relevant for studying obesity-independent NAFLD. Commonly used high-fat diet-induced NAFLD animal models are not ideal for testing this hypothesis.

**APPROACH AND RESULTS:** In this study we used a transgenic zebrafish with chronic systemic overexpression of human IL6 (IL6-OE) and found accumulation of triglyceride in the liver. We performed comparative transcriptomics and proteomics on the IL6-OE liver and found an expression signature distinct from the diet-based NAFLD models. We discovered a deregulation of glycolysis/gluconeogenesis pathway, especially a robust down regulation of the glycolytic enzyme *aldolase b* in the IL6-OE liver. Metabolomics of the IL6-OE liver showed accumulation of hexose monophosphates and their derivatives, which can act as precursors for triglyceride synthesis. Patients with the genetic disease Hereditary Fructose Intolerance (HFI) caused by *ALDOLASE B* deficiency also have a higher propensity to develop fatty liver disease.

**CONCLUSIONS:** Our study demonstrates a causative role for inflammation in intrahepatic lipid accumulation. Further, our results suggest that IL6-driven repression of glycolysis/gluconeogenesis, specifically *aldolase b*, may be a novel mechanism for development of fatty liver, especially in obesity-independent NAFLD.

---

Non-alcoholic fatty liver disease (NAFLD) has become a major public health concern worldwide with an estimated 20-30 % of total population affected by NAFLD (1, 2). NAFLD is a chronic liver disease, which begins with steatosis or accumulation of fat in the liver, progresses to non-alcoholic steatohepatitis (NASH) and eventually develops into fibrosis, cirrhosis, liver cancer and finally liver failure (3). It is strongly associated with obesity, dyslipidemia, and metabolic syndrome (4), however recently NAFLD has been noted in lean individuals, especially in Indians (5, 6). These observations suggest that there might be independent mechanisms for developing fatty liver.

Inflammation finds constant mention in studies of intrahepatic lipid accumulation. For instance, exogenous treatment of HepG2 cells with palmitic acid or oleic acid promotes the accumulation of fat and induces pro-inflammatory cytokines such as IL8 (7) and TNFalpha (8) respectively. Rats fed on a high-fat diet (HFD) develop steatosis in the liver eventually leading to steatohepatitis (9). The extent of steatosis in NAFLD patients is strongly correlated with the expression of pro-inflammatory cytokines such as TNFalpha, IL6 and IL1beta (10-12). So far, most studies have suggested that steatosis precedes inflammation in the liver. However, there are indications for chronic inflammation playing a more determinant role in fatty liver. Recent studies found that *Klebsiella pneumoniae* induces high IL6 expression in gut and liver and also causes hepatic steatosis (13). Patients of chronic inflammatory diseases such as Systemic Lupus Erythematosus (SLE), Crohn’s disease, Inflammatory Bowel Disease (IBD) and Rheumatoid Arthritis (RA), all of whom have high systemic IL6 (14-17), appear to develop steatosis (18-21). Treatment of HepG2 cells with pro-inflammatory cytokines IL1beta and TNFalpha also promotes steatosis (22, 23).

Most available models of NAFLD are fed on high-fat diet (HFD). To study the relationship between chronic inflammation and steatosis in zebrafish here we use a Gal4-UAS system that drives constitutive expression of human IL6 in the heart. Chronic exposure to secreted IL6 induced steatosis in the adult male liver. We characterized the gene expression changes in the IL6-transgenic zebrafish liver by RNA sequencing and proteomics studies. The classic NAFLD expression signatures were absent in the IL6-transgenic liver; instead we found a down regulation of the glycolytic/gluconeogenesis pathway. Specifically, we discovered a repression of the expression of *aldolase b* and a resulting accumulation of C6-monophosphates. We propose that chronic inflammation leads to fatty liver through deregulation of glycolytic flux. This model with chronic systemic-inflammation-induced steatosis in zebrafish would be valuable in understanding the causative role of inflammation in fatty liver disease and would also be useful for designing chemical screens for this flavor of NAFLD.

## Materials and Methods

Additional information is provided in the Supporting Materials and Methods.

### ZEBRAFISH LINES AND MAINTENANCE

Zebrafish (*Danio rerio*) were bred, raised, and maintained at 28.5□°C under standard conditions as previously described (24). Zebrafish handling was in strict accordance with good animal practices as described by the Committee for the Purpose of Control and Supervision of Experiments on Animals (CPCSEA), Government of India. The Institutional Animal Ethics Committee (IAEC) of the CSIR-Institute of Genomics and Integrative Biology, New Delhi, India approved all animal experiments. The transgenic zebrafish lines used in this study are Tg(pCH-cmlc2:GVP), Tg(pCH-cmlc2:GVP::pBH-UAS:IL6) (25), Tg(pCH-fabp10a:GAL4m:: pBH-UAS:IL6) (26).

### RNA IN SITU HYBRIDIZATION

The embryo (96 hours post fertilization) and heart tissues from adult zebrafish were harvested and fixed in 4% ice cold paraformaldehyde overnight at 4°C. RNA in situ hybridization was performed as previously described (24). Digoxigenin-labelled antisense riboprobe against human IL6 gene was used in this study.

### QUANTITATIVE REAL TIME PCR

Total RNA was extracted from zebrafish embryos, adult heart, liver or HepG2 cells, using RNAiso Plus (Takara, 9109) reagent. cDNA synthesis was done using a QuantiTect Reverse Transcription kit (Qiagen, 205313). Quantitative real time PCR was performed as previously described (27) using SYBR green (Roche, 06924204001) reagent on ROCHE LightCycler 480 System. We used relative standard curve method for quantification of RNA transcript as described by the manufacturer to generate raw values in arbitrary units. Each experiment was performed with at least three replicates. The normalized data were analyzed using 2-ΔΔCT method. All quantifications were normalized to rpl13α unless mentioned otherwise. For statistical significance of the data, p-values were calculated by performing unpaired Student t test. The GraphPad Prism 5.04 software was used for plotting the graphs. All primers used in this study are listed in Supporting Table S1.

### WESTERN BLOT ANALYSIS

Adult zebrafish liver and heart tissue were homogenized in NP40 lysis buffer (ThermoFisher, FNN0021). Protein concentrations were estimated using BCA (Pierce 23225) method and equal amount of protein (25µg) was loaded in all well. Western blots were performed as described (28). Primary antibodies against humanIL6 (abcam-ab6672), Aldolase B (GeneTex-GTX124306), STAT3 (Cell Signaling-4904s), phospho-STAT3 (Cell Signaling-9131s) were used at 1:1000 dilution. Secondary antibody was HRP conjugated and the signal was detected using EMD Millipore Immobilon Western Chemiluminescent HRP Substrate (Merch-WBKLS0500) on a G:BOX (Syngene) detection system.

### HISTOLOGY AND STAINING

Zebrafish liver tissues were fixed in 4% paraformaldehyde. Sections were 7µm thick. Cryo-sections were used for Oil Red O and BODIPY (493/503) staining and paraffin embedded sections were used for Hematoxylin and Eosin (H&E) staining. BODIPY sections were mounted with DAPI-containing mounting media. Nikon upright microscope (model E200) was used for Oil Red O and H&E sections. Leica SP8 confocal microscopy was used for BODIPY stained sections.

### TRANSMISSION ELECTRON MICROSCOPY

Adult zebrafish liver was fixed in 2.5% glutaraldehyde and 4% paraformaldehyde. The section were osmicated in 1% osmium tetroxide followed by dehydration in graded series of alcohol, followed by infiltration with Epon 812 resin. Ultra-thin sections were cut (60-70 nm) on Ultramicrotome (Leica) and placed on copper grids. The sections were stained with uranyl acetate and lead citrate. Finally, the samples were visualized on a Transmission Electron Microscope (Tecnai G2 20 twin, FEI).

### LIPID EXTRACTION AND THIN LAYER CHROMATOGRAPHY

Total lipids were extracted from liver tissue using Modified Bligh and Dyer Method. The dried lipid extracts re-suspended in chloroform:methanol (2:1) were loaded on thin-layer chromatography plate and separated in the solvent system hexane:diethyl ether:acetic acid (70:30:1), used for neutral lipids. The plates were stained using 10% copper sulfate (w/v) in 8% phosphoric acid (v/v) solution followed by charring at 150°C. Quantification of un-labeled lipid spots was done using ImageJ® software (29).

### TRANSCRIPTOMICS

RNA was isolated from adult liver (from 3 animals in each group) of control males and females and IL6-OE males and females. RNA-seq libraries were prepared from 1µg of total RNA using standard protocol. RNA sequencing was performed on HiSeq2500 sequencing platform from Illumina, USA. Details of sequencing and preliminary analysis protocols are in Supplementary Methods.

### DATA DEPOSITION

The raw reads file in Fastq format has been uploaded on the NCBI Short Read Archive (SRA). The data can be accessed on NCBI Bioproject: PRJNA638724. The list of differentially expressed genes is available in Supporting Table S2.

### PROTEOMICS

Adult liver protein from control males and females and IL6-OE males and females (2 animals of each group) was extracted and three replicate 8-plex experiments were performed. The proteins were labeled using 8plex iTRAQ reagent as per the manufacturer’s protocol (AB Sciex). The peptides were separated on Eksigent nano-LC (Ultra 2D) coupled with 5600 triple time-of-flight (TOF) (AB Sciex). Details of the proteomics experiment and preliminary analysis protocols are in Supplementary Methods. The list of differentially expressed proteins is available in the Supporting Table S3.

### METABOLOMICS ANALYSIS OF HYDROPHILIC METABOLITES

Liver samples were cryo-fixed and lyophilized at −100°C for 24 hours in cryo-lyophilizer (Benchtop Pro with omnitronics, SP scientific) and stored at −80 °C till further extraction. The complete details for sample extraction for hydrophilic metabolites, analysis by an orbitrap mass spectrometer and data analysis is given in Supplementary Methods section. The median values for the metabolites from 8 replicate samples (4 biological and 4 technical) were used to calculate the fold change values and identify the metabolites showing significant change (≥ & ≤ 2 fold change; p-value ≤ 0.05). The raw and processed metabolite data have been deposited in the metabolomics workbench repository.The list of differentially expressed metabolites is available in the Supporting Table S4.

### CELL CULTURE

HepG2 cells were cultured at semi-confluent densities for the experiments. Cells were treated with recombinant IL6 (abcam-ab119444). All experiments were performed at least in triplicates.

## Results

### OVEREXPRESSION OF HUMAN IL6 IN THE ZEBRAFISH HEART INDUCED IL6 SIGNALING IN THE LIVER

To probe the role of chronic systemic inflammation on hepatic steatosis, we overexpressed the pro-inflammatory cytokine, human interleukin 6 (Hsa.IL6; henceforth called ‘IL6’) in the zebrafish heart. We used a double transgenic system *Tg (myl7:GAL4-VP16)::(UAS:HsaIL6)* (25). Here the myl7 promoter drives GAL4-VP16 expression in the heart, which induces specific overexpression of IL6 (Fig. 1A). RNA in situ hybridization of 4 day old zebrafish embryos with a human IL6 specific probe indicated strong expression in the embryonic heart in the double transgenic line (henceforth IL6-OE) but not in the control embryos of *Tg (myl7:GAL4-VP16)* (Fig. 1B).

**Figure 1.**
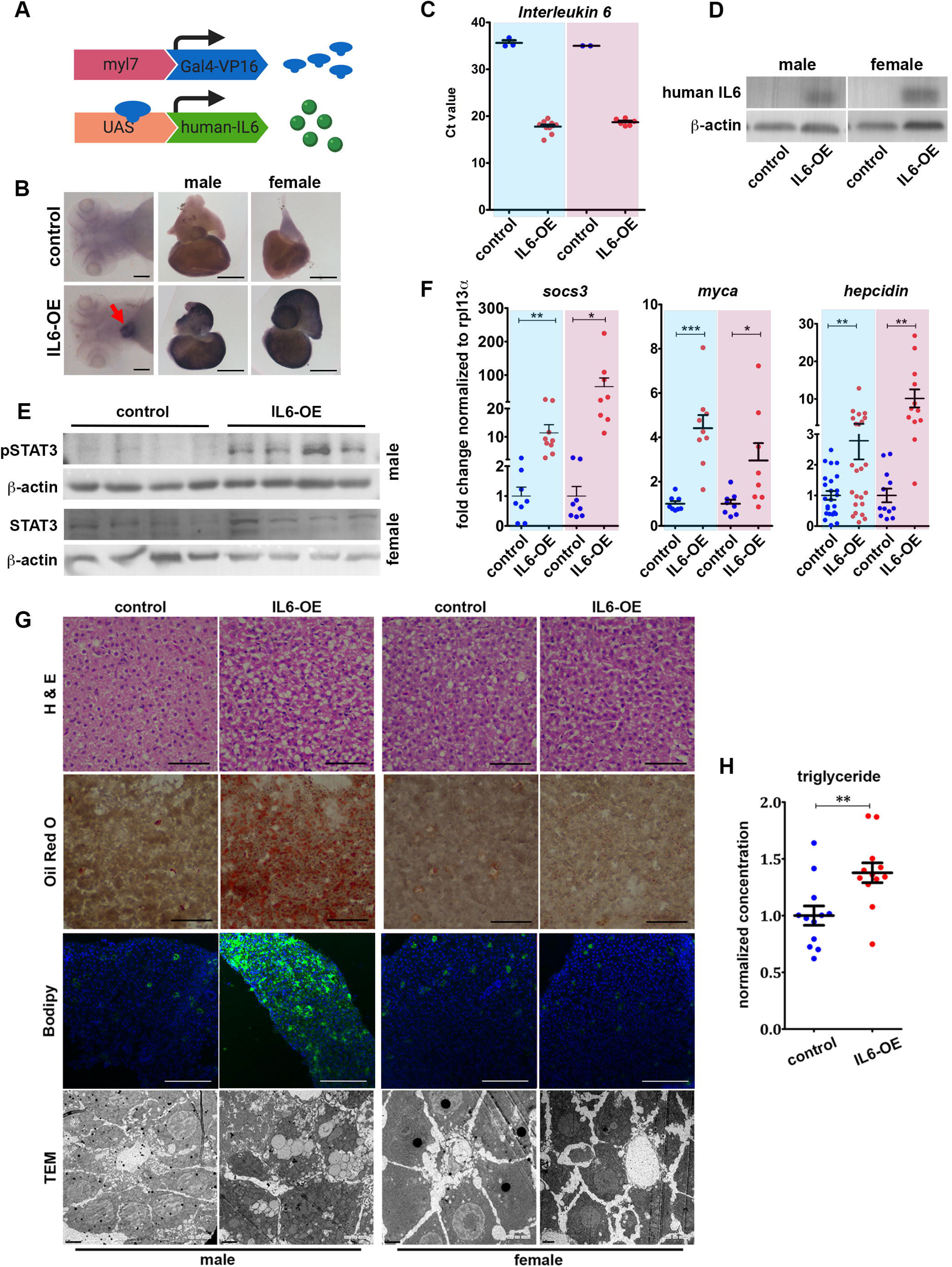
Over-expression of human IL6 in zebrafish heart induces lipid accumulation in the liver. (A) Schematic depiction of the double transgenic *Tg(pCH-cmlc2:GVP::pBH-UAS:IL6)* which leads to expression of human IL6 constitutively in the zebrafish heart. (B) RNA in situ hybridization on 4 day old zebrafish embryos and adult heart shows expression of human IL6 in the heart (red arrow in embryo) in IL6-OE. Embryos are presented in ventral view, anterior to the left. Scale bar 100µm. Adult heart shows atrium to the top and ventricle to the bottom. Scale bar 500µm. (C) Ct values from qRT-PCR of human IL6 on adult heart tissue shows low Ct values i.e. high expression in the IL6-OE samples. Data in blue box is for male and pink box is for female. (D) Western blot of adult heart shows a 21kDa IL6 band in the IL6-OE male and female. Beta-actin is used as loading control. (E) Western blot of adult liver shows p-STAT3 induction in the IL6-OE, strongly in male than in female. (F) qRT-PCR quantification of STAT3 target genes, *socs3, myca* and *hepcidin* in the liver of IL6-OE fish. Data in blue box is for male and pink box is for female. The data is fold change normalized to rpl11alpha is presented as mean ± SE. (G) Histological examination of liver sections of male and female control and IL6-OE animals. Hematoxylin and Eosin (H&E) reveals distinct morphology of male IL6-OE sample. Large white spaces pushing hepatocyte nuclei to the periphery compared to packed centrally nucleated hepatocytes in control. Female control and IL6-OE are similar. Oil red O staining shows lipid accumulation in male IL6-OE. No significant lipid presence in the other samples. BODIPY 493/503 shows lipid accumulation (green) in male IL6-OE. Sections are counterstained with phalloidin (red) and DAPI (blue). Scale bar 50µm (H&E, oil red O and BODIPY). TEM images show ultrastructure of liver. Large lipid droplets are visible in the male IL6-OE. Scale bar 2µm. (H) relative fold change of triglyceride in male IL6-OE liver by thin layer chromatography (Supplementary Fig. 2D). Data presented mean ± SE. *p* value is calculated using unpaired student t-test, * p< 0.05, ** p< 0.01 and *** p< 0.001. Each point in graph represents a single animal. Abbreviations: H&E, Hematoxylin and eosin; TEM, Transmission electron microscopy.

We performed RNA in situ hybridization on adult heart tissue of IL6-OE and control animals with the human IL6 probe and found strong expression in the heart of both male and female IL6-OE animals compared to controls (Fig. 1B). Quantitative RT-PCR (qRT-PCR) of adult hearts was performed and the raw Ct values (corrected to rpl13α) were plotted. There was a difference of nearly 20 cycles between the control and IL6-OE hearts (Fig. 1C). Western blotting found a 21 kDa band corresponding to the human IL6 in the adult hearts of IL6-OE animals with no signal in the control animals (Fig. 1D). Western blotting for phosphorylated STAT3 in the adult liver of control and IL6-OE animals showed the induction of an 88 kDa band for p-STAT3 in the IL6-OE liver (Fig. 1E) suggesting IL6 signaling in the liver. Quantification of p-STAT3 showed an average induction of 3.47 fold in male IL6-OE livers; the induction was not significant in female (Supplementary Fig. 1A-B). Quantification of known target genes of IL6 such as *socs3* (30) showed an induction of 11.4 fold in male and 65.4 fold in female IL6-OE (Fig. 1F). IL6 target genes *myca* and *hepcidin (HAMP*) were also strongly induced in male and female IL6-OE adult liver (Fig. 1F). These experiments indicated that human IL6 was expressed specifically in the heart and secreted into the plasma leading to IL6 signaling in the liver.

### CHRONIC OVEREXPRESSION OF IL6 INDUCES HEPATIC STEATOSIS IN IL6-OE MALE ZEBRAFISH

Dissection of adult livers from IL6-OE animals revealed an enlarged and discolored appearance in the male tissue (Supplementary Fig. 2A). There was a significant increase in dry weight of IL6-OE liver in male and female animals compared to the respective controls (Supplementary Fig. 2B). We performed Hematoxylin and Eosin staining on tissue sections and found that the male IL6-OE liver had a distinctive appearance compared to the control livers. In the control animals and the female IL6-OE, the hepatocytes were tightly packed with central nuclei while in the IL6-OE livers there were prominent white vesicles/spaces that appeared to push the hepatocyte nuclei to the side (Fig. 1G). Since the appearance of the male IL6-OE liver sections was reminiscent of fatty liver, we performed Oil red O staining. The adult liver in control and female IL6-OE had the presence of a few red droplets indicating lipid (Fig. 1G).

However, the male IL6-OE liver had more lipid droplets at 8 months old (Supplementary Fig. 2C) and at 2 years old the section was filled with lipid (Fig. 1G). Staining with the fluorescent dye BODIPY 493/503 to detect lipid showed that the male IL6-OE liver had higher lipid staining than the control male, control female, and the IL6-OE female (Fig. 1G). We performed transmission electron microscopy to look at the ultrastructure of hepatocytes in the adult liver and found that compared to control male, the IL6-OE male had hepatocytes filled with large lipid droplets (Fig. 1G). The female control and IL6-OE livers had a very similar but distinct-from-male ultrastructure (Fig. 1G). We performed thin layer chromatography to identify and quantify the types of lipids in the male IL6-OE liver. Compared to standards run alongside we found a 1.3 fold increase in triglycerides and a 1.2 fold increase in cholesterol/diacylglycerol in the male IL6-OE liver compared to the control male liver (Fig. 1H, Supplementary Fig. 2D-E). Serum measurements showed a decrease in triglycerides and glucose but no change in cholesterol in IL6-OE male animals (Supplementary Fig. 2F-H).

### CHRONIC EXPOSURE TO IL6 SUPPRESSES LIPOGENESIS, B-OXIDATION AND AUTOPHAGY IN IL6-OE MALE LIVER

Lipid accumulation in hepatocytes could result from increased lipogenesis or decreased lipolysis in the liver. Previous studies on HFD models have shown increased fatty acid synthesis (31) and induction of FASN expression (32) in NAFLD and hepatic steatosis patients, respectively. A fatty liver model induced by acute inflammation using intraperitoneal injections of TNFalpha in mice also reported induction of fatty acid synthase (23). Contrary to observations in diet-induced NAFLD models, we found a statistically significant down regulation of *acaca, srebp1, pparab, pparg* and *fads2* in the IL6-OE livers by qRT-PCR (Fig. 2A).

**Figure 2.**
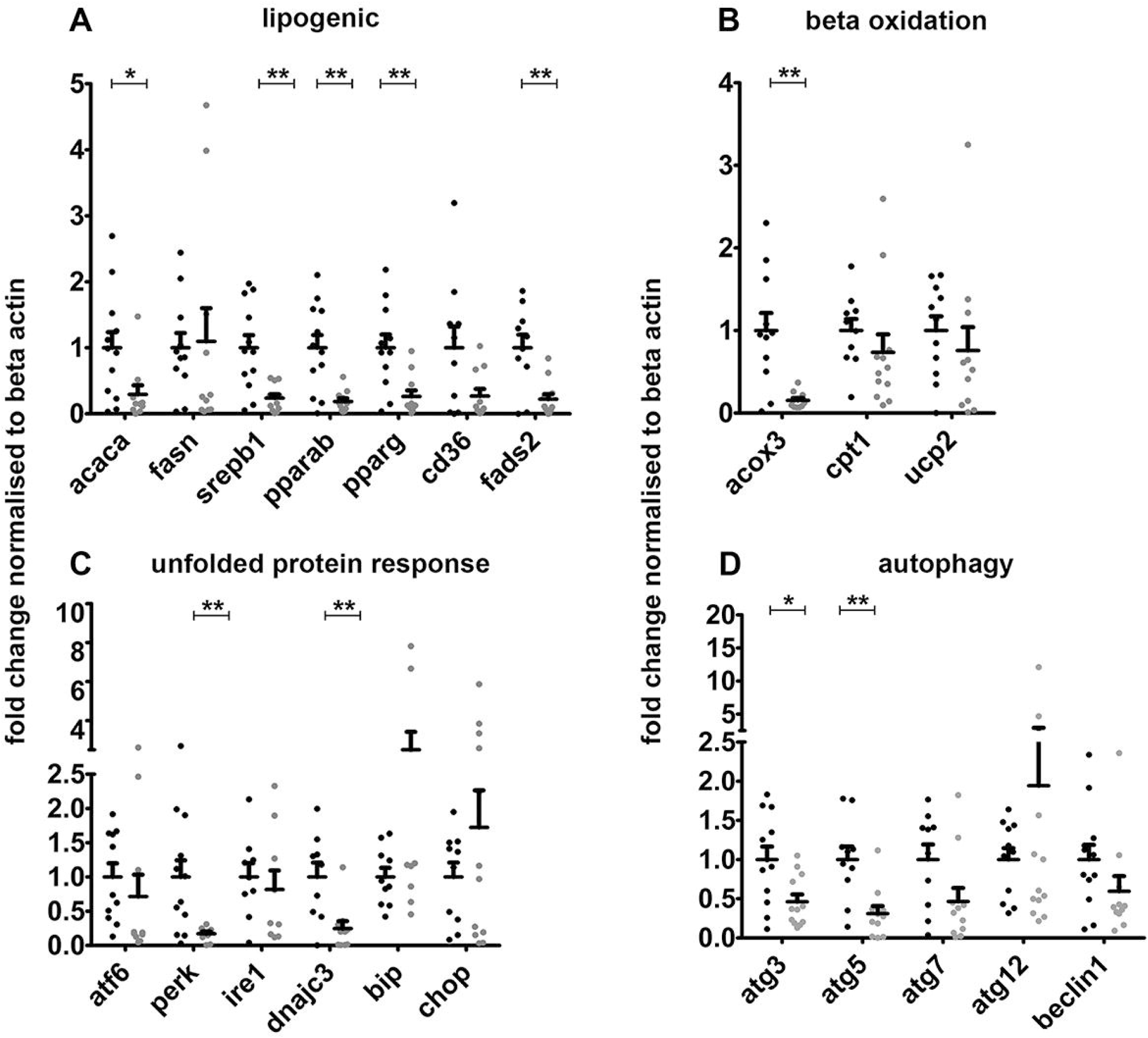
Chronic exposure to IL6 suppresses lipogenesis, β-oxidation and autophagy in male liver. qRT-PCR of genes known to be regulated in NAFLD. Fold change normalized to beta-actin is plotted. (A) Expression of genes involved in lipogenesis. Significant down regulation was observed in *acaca, srebp1, pparab, pparg* and *fads2* in IL6-OE. (B) Expression of the beta-oxidation pathway genes. Peroxisomal beta-oxidation gene *acox3* is down regulated in IL6-OE. (C) Expression of genes involved in unfolded protein response pathway. Significant down regulation of *perk* and *dnajc3* was seen in IL6-OE. (D) Genes of autophagy pathway, atg3 and atg5 were significantly down regulated in IL6-OE. The data is presented as mean ± SE. *p* value is calculated using unpaired student t-test, * p< 0.05 and ** p< 0.01. Each point in graph represents single animal.

Unlike diet-based mouse models, NAFLD patients show negative correlation between hepatic *PPAR alpha* expression and severity of steatosis/presence of NASH (33). The down regulation of PPAR signaling in the zebrafish IL6-OE suggests that it might serve as an alternative to model NAFLD in humans. In a recently published study adult male zebrafish exposed to bisphenol S accumulate fat in the liver (34). This model also shows a down regulation of *PPAR alpha* expression. The progression of NAFLD is associated with suppression of fatty acid oxidation (35). Comparison of expression of the mitochondrial beta-oxidation genes *cpt1* and *ucp2* showed no significant change in IL6-OE livers. The expression of *acox3*, involved in peroxisomal beta-oxidation, was significantly down regulated in the IL6-OE male liver (Fig. 2B).

Lipid accumulation in NAFLD and NASH induces endoplasmic reticulum stress in the liver (36). We quantified the expression of unfolded protein response (UPR) genes in the IL6-OE male liver and found instead, a down regulation of *perk* and *dnajc3* and no change in *aft6, ire1, bip*, and *chop* (Fig. 2C). Another pathway that is up regulated in NAFLD is autophagy (37). In IL6-OE male liver the levels of autophagy genes *atg3* and *atg5* were significantly down regulated, while *atg7*, and *beclin*-1 showed a downward trend (Fig. 2D). We quantified the expression of genes involved in dietary fat absorption in gut (Supplementary Fig. 3A-B), lipid transport in the liver (Supplementary Fig. 3C-E) and bile acid metabolism in liver (Supplementary Fig. 3F-K) using qRT-PCR. We found no change in any of these genes except a significant up regulation of *abca1b* (lipid export), *cyp7b1* (bile acid metabolism) and down regulation of *slc27a2* (lipid import) (Supplementary Fig. 3C-D, I), which was counter intuitive.

### DISTINCT TRANSCRIPTOMIC SIGNATURES IN THE IL6-OE ZEBRAFISH LIVER

Since the IL6-OE zebrafish liver appeared to not have the usual transcriptomic patterns, we decided to compare the whole transcriptome of the control and IL6-OE liver of male and female by RNA sequencing (each n=3). We performed the RNA sequencing using Illumina Hiseq 2500 platform. To assess the transcriptomic concordance between samples we computed the Spearman correlation between the gene expression profiles. We found that the sample ‘female IL6-OE-3’ did not cluster with the other two samples of the group and this sample was removed from further analysis (Supplementary Fig. 4A). We obtained an average depth of 52 million reads and mapped the reads to the zebrafish reference genome (Zv11) achieving >60-89% alignment using Bowtie tool (38). The transcripts were annotated using the reference transcript of ENSEMBL v87 and FPKM of each gene from all the samples were calculated using Cuffnorm (39). We removed all genes with an FPKM=0 in any of the samples to simplify the analysis. The FPKM of genes from IL6-OE male and female livers were compared to their respective controls. 1755 genes with a minimum of 2 fold change difference between control and IL6-OE at a statistical significance of p<0.05 by Student t-test were shortlisted for the final differentially expressed gene (DEG) list (Fig. 3A). Clustering of the genes based on their expression pattern showed distinct groups of genes that are up regulated and down regulated in the male and female livers. A Venn diagram of the 1755 DEGs in our dataset showed that 436 genes were up regulated and 508 were down regulated in the male IL6-OE liver compared to the control male liver (Fig. 3B). Of these, 42 up regulated genes and 103 down regulated genes were common to both male and female IL6-OE liver. Since the fatty liver phenotype was seen only in the male IL6-OE liver, we performed all further analysis on the 791 DEGs that were unique to the male IL6-OE liver.

We performed a gene ontology analysis using DAVID database (40) on the 791 DEGs for enrichment of pathways in the transcriptome. Among the genes up regulated, one of the most prominent class was the JAK-STAT signaling pathway (Supplementary Fig. 4B), confirming the systemic effect of IL6 protein expression from the heart. The other striking group of genes was those involved in protein processing in ER (Supplementary Fig. 4B, 5A). The Unfolded Protein Response genes we had shown to be down regulated by qRT-PCR (Fig. 2C) were distinct from the ones present in the DEGs (Supplementary Fig. 5A).

Among the genes down regulated in male IL6-OE, metabolism was the dominant class (Fig. 3C). In accordance with our qRT-PCR results that showed down regulation of lipid metabolism genes (Fig. 2A-B) we found a partially overlapping set of genes down regulated in the RNA sequencing data (Supplementary Fig. 5B). The transcriptomics data confirmed the down regulation of genes involved in PPAR signaling (Supplementary Fig. 5C) as observed in our earlier experiments (Fig. 2A). An interesting class of genes down regulated exclusively in the male and not in the female IL6-OE liver was the glycolysis/gluconeogenesis pathway (Fig. 3C). Of the glycolysis genes (Supplementary Fig. 4C), we found that 9 were dramatically down regulated in IL6-OE male liver (Fig. 3D). qRT-PCR of some of these genes e.g. *g6pca, pgm1, pgm2, pgam2* and *pdhb* validated the RNA sequencing profile (Supplementary Fig. 6A-E).

To strengthen our findings, we carried out global protein profiling of the liver from control and IL6-OE male and female zebrafish (each n=2) in three replicate by Liquid Chromatography Tandem Mass Spectrometry (LC-MS/MS) using iTRAQ labeling. We shortlisted only those proteins for analysis that were represented by at least 2 peptides, with fold change <=0.8 or >=1.2. The data was analyzed using protein pilot (SCIEX) and a total of 138 significantly altered proteins were identified in all the groups. 16 proteins were up regulated only in male and 95 proteins only in female IL6-OE. 4 proteins were uniquely down regulated in the male and 11 in female IL6-OE (Fig. 3E). DAVID gene ontology analysis showed an enrichment of the protein processing pathway in ER in both male and female IL6-OE zebrafish and the ribosomal machinery only in the female IL6-OE (Supplementary Fig 6F). Carbon metabolism was enriched exclusively in the male IL6-OE liver with down regulation of four proteins: Aldob, Hemopexin, Herc4 and Ethanolamine phosphate phospholyase (Supporting Table S3).

Aldolase b (Aldob), an important enzyme in the glycolysis pathway (Supplementary Fig. 4C), was the only gene that showed a consistent down regulation in the male IL6-OE liver, both at the RNA (Supplementary Fig 7A) and protein level (Supplementary Fig 7B). qRT-PCR with a larger sample set showed a 1.53 fold down regulation of *aldob* RNA and western blot analysis showed a 1.68 fold down regulation in Aldob protein levels (Fig. 3F-G, Supplementary Fig. 7D). Since the effect seen on the liver is due to IL6 secreted from heart we used a Gal4-UAS based transgenic line, *Tg(fabp10a:GAL4m::pBH-UAS:IL6)* (26), which expresses human IL6 specifically in the hepatocytes (henceforth known as hepIL6-OE). By western blot we found a significant induction of p-STAT3 in the hepIL6-OE liver indicating robust IL6 signaling (Supplementary Fig. 7E-F). We then compared the Aldob protein levels in the same liver and found a 2.95 fold down regulation of in hepIL6-OE compared to controls (Supplementary Fig. 7E,G).

### CHRONIC IL6 EXPOSURE PROMOTES INTRAHEPATIC ACCUMULATION OF HEXOSE MONOPHOSPHATE IN MALE ZEBRAFISH

The RNA sequencing and LC-MS data showed a down regulation of the glycolysis gene *aldob*. Aldolase b enzyme [EC 4.1.2.13] converts fructose-1, 6-bisphosphate into glyceraldehyde-3-phosphate and dihydroxyacetone phosphate (DHAP) (Supplementary Fig. 4C). We performed untargeted metabolomics for polar metabolites in the IL6-OE zebrafish liver using LC-MS to detect any metabolic perturbation (each n=4). We compared the levels of 120 metabolites and found 32 metabolites to be differentially accumulated at a set threshold of 2-fold change and p<0.05 in the male IL6-OE liver (Supplementary Fig. 8B). We found a 1.55 fold accumulation of beta-D-fructose-1,6 bisphosphate, the substrate of Aldolase b, in the IL6-OE male liver (Supplementary Fig. 8C). There was also a 4.60 fold accumulation of hexose-monophosphates in the IL6-OE liver (Fig. 3H). Studies on patients who are genetically ALDOB deficient have suggested intrahepatic accumulation of hexose monophosphate (42) suggesting a similar change in metabolic flux as the disease Hereditary Fructose Intolerance. Various other sugar-phosphate forms, which are part of the pentose phosphate pathway also accumulated in the male IL6-OE liver (Supplementary Fig. 8B).

One of the highest change was that of DHAP/glyceraldehyde-3-phosphate (these two molecules cannot be resolved on the LC-MS, Fig. 3I), which showed a 7.03 fold accumulation (Fig. 3I). Our transcriptomics data shows a repression of *gapdh*, which uses Glyceraldehyde 3-phosphate as a substrate (Fig. 3D). Further, our proteomics data revealed an up regulation of Tpi1b, the isomerase that interconverts DHAP and glyceraldehyde 3-phosphate (Supplementary Fig. 7C) suggests a favored conversion of glyceraldehyde 3-phosphate to DHAP. DHAP is further converted to Glycerol 3-phosphate by the enzyme Gpd2. We observed a 2.48 fold accumulation of sn-glycerol-3-phosphate, although not statistically significant, (Supplementary Fig. 8D) in the male IL6-OE liver and a 2.14 fold induction in the mRNA coding for *gpd2* in transcriptomic study (Fig. 3J). DHAP is a substrate in the triacyl glycerol (triglyceride) synthesis pathway with glycerol-3-phosphate as an intermediate suggesting the mechanism for triglyceride accumulation in the male IL6-OE liver (Fig. 3K and Supplementary Fig. 4C).

### IL6 TREATMENT INHIBITS ALDOB EXPRESSION IN HEPG2 CELLS

To test the causative relationship between IL6 and ALDOB expression, we treated the human liver carcinoma cell line HepG2, with recombinant human IL6. To identify the optimum concentration of rIL6 we treated HepG2 cells with different concentrations of rIL6 for 24 hours. We found a dramatic reduction in ALDOB protein levels at 10ng/mL, which did not change at higher concentrations (Fig. 4A). To study the kinetics of repression, we treated HepG2 cells with 100ng/ml rIL6 for different time periods and measured the *ALDOB* mRNA levels. We found a strong statistically significant repression of *ALDOB* at 24 hours (Fig. 4B). Finally a treatment with 100ng/mL IL6 for 24 hours caused a 2.07 fold down regulation of *ALDOB* mRNA expression across multiple experiments (Fig. 4C).

**Figure 3.**
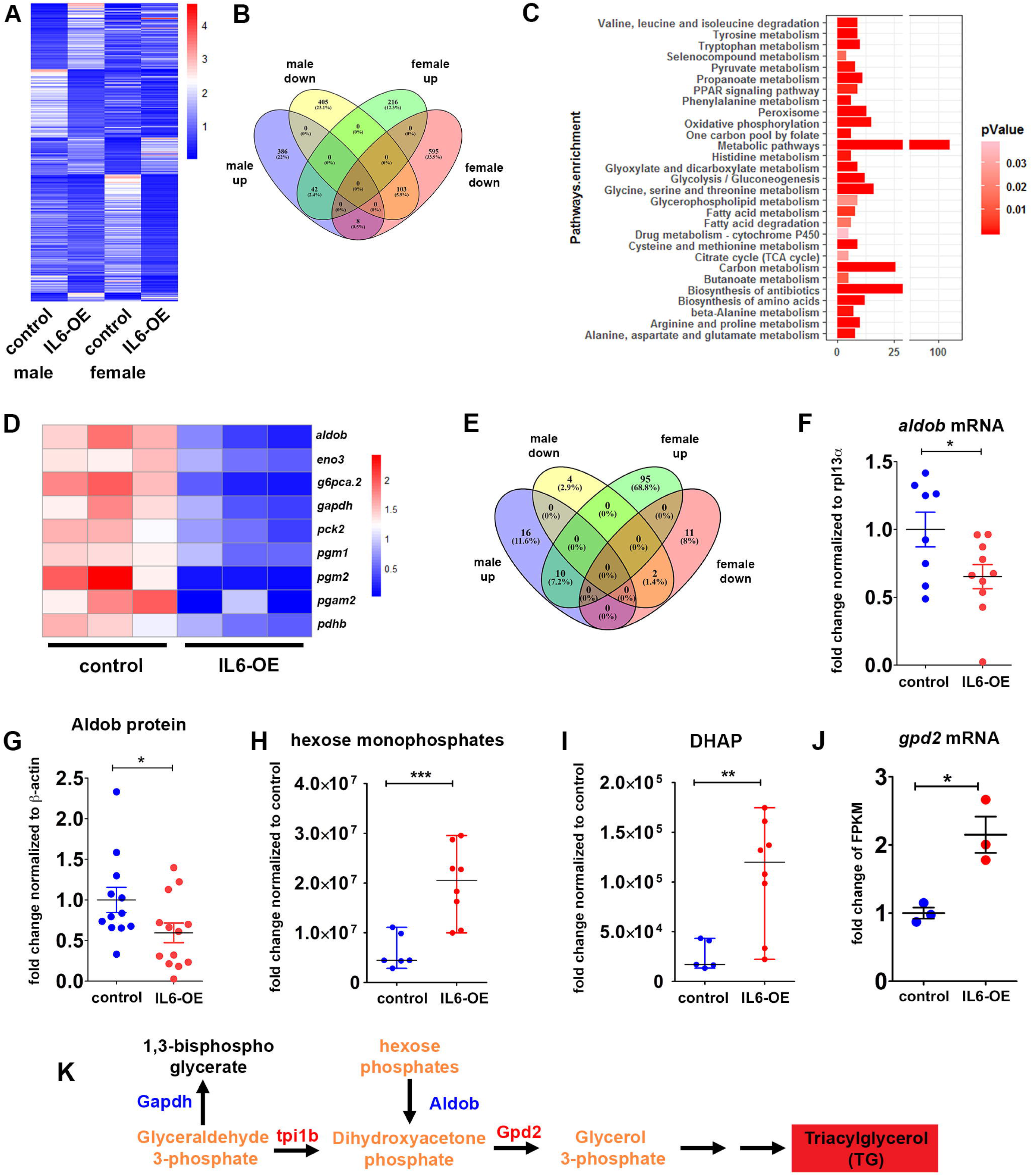
Male IL6-OE zebrafish liver exhibits a distinct expression profile with down regulation of glycolysis/gluconeogenesis pathway. (A) 1755 genes were differentially expressed in the IL6-OE compared to the respective controls. Average of three samples in each group is depicted. FPKM of each group is normalized to FPKM average of the respective gene. (B) 791 genes were uniquely differentially expressed in the male IL6-OE liver. (C) Pathways enrichment analysis of genes down regulated in the male IL6-OE using DAVID algorithm shows large metabolic changes. (D) 9 genes encoding glycolysis enzymes are down regulated exclusively in the male IL6-OE liver. FPKM of each sample is normalized to FPKM average of the respective gene. (E) Proteomics studies revealed that 138 proteins were differentially expressed in the IL6-OE zebrafish liver. (F-G) qRT-PCR of adult male liver showed a 1.53 fold down regulation of *aldob* mRNA and Western blot quantification showed a 1.68 fold down regulation in Aldob protein levels in the male IL6-OE liver. (H-I) Metabolic profiling revealed accumulation of Hexose-monophosphate and DHAP in the IL6-OE liver. (J) Expression of *gpd2* transcript which converts DHAP into glycerol-3-phosphate were significantly up regulated in IL6-OE male liver. (K) Proposed model of conversion of DHAP to triglyceride accumulation in IL6-OE liver. (F-G, J) The data is presented as mean ± SE and (H-I) The data is presented as median ± Range. *p* value is calculated using unpaired student t-test, * p< 0.05. Each point in graph represents single animal.

**Figure 4.**
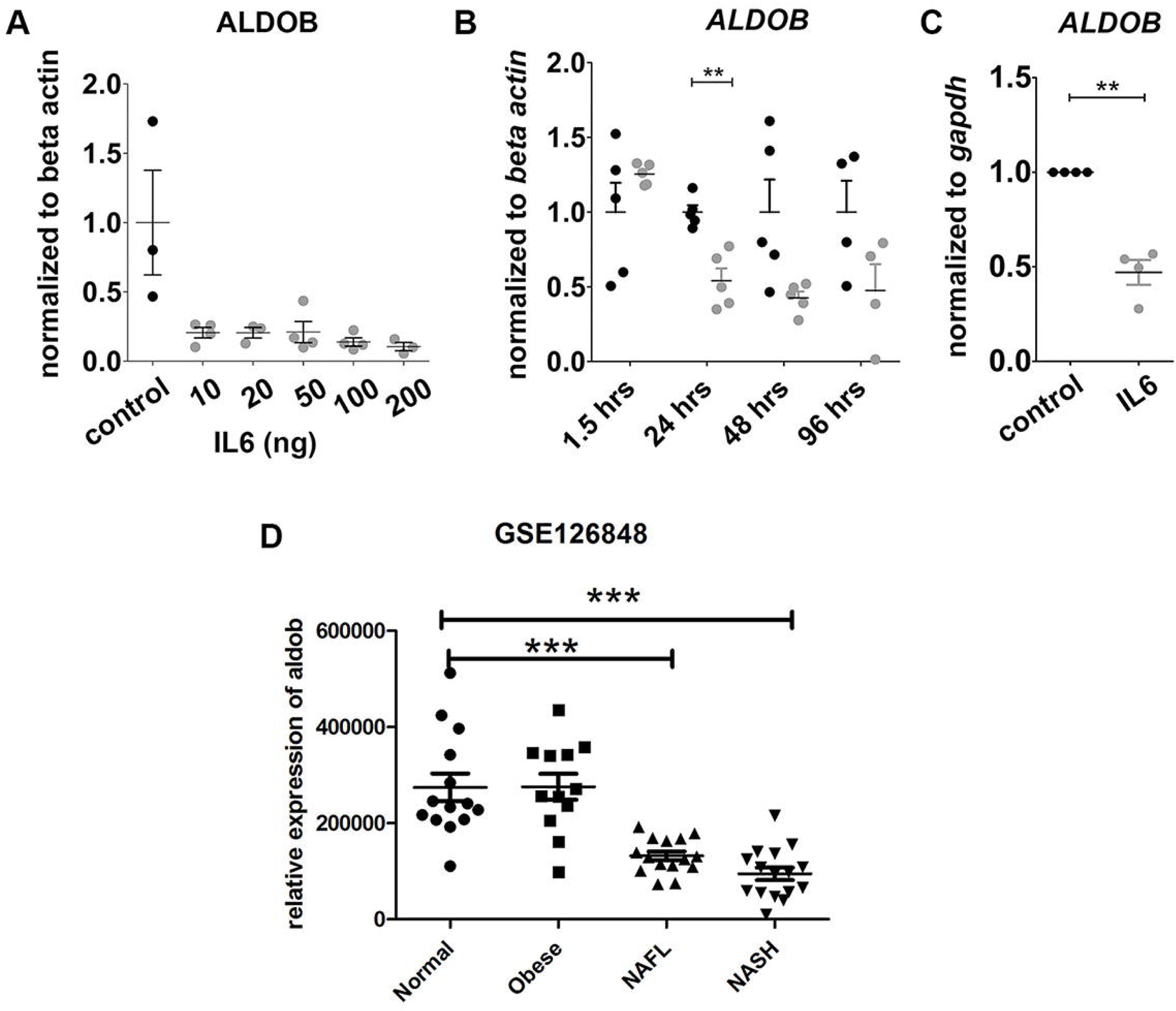
Lower ALDOB levels in IL6 treated cells and NAFLD patients. (A) HepG2 cells treated with increasing concentrations of recombinant human IL6 for 24 hours show robust repression of ALDOB protein. (B) Treatment with 100ng/ml of rIL6 for different duration shows minimum *ALDOB* mRNA levels at 24 hours. (A-B) The data is presented as mean ± SE. *p* value is calculated using unpaired student t-test, ** p< 0.01. Each point in graph represents single sample. (C) Exposure of HepG2 cells to 100ng/ml of rIL6 for 24 hours shows a robust repression of *ALDOB* transcript. The data is presented as mean ± SE. *p* value is calculated using unpaired student t-test, ** p< 0.01. Each point in graph represents one experiment with each experiment conducted on three samples each. (D) Analysis of relative expression levels of *ALDOB* in publicly available dataset GSE126848 reveals a 2.08 fold reduction in NAFLD and 2.9 fold down regulation in NASH patients. There is no significant difference between normal and obese individuals. The data is presented as mean ±SE. *p* value is calculated using unpaired student t-test, *** p< 0.001.

### ALDOB EXPRESSION CORRELATES WITH NAFLD IN PATIENTS

Our studies show that chronic systemic IL6 expression leads to a repression of aldob in the liver and accumulation of hexose monophosphates. ALDOB deficient individuals have been reported to have high intrahepatic triglyceride content (41). We hypothesized that ALDOB levels and the resulting metabolic changes would have a correlation with fat accumulation in patients. To test this hypothesis we looked for gene expression profiles of NAFLD patients in the public domain. We found a previous study that had generated transcriptomics profile for liver from normal, obese, NAFLD and NASH human subjects to improve diagnostic criteria in patients (42). This dataset was publically available (GEO accession number-GSE126848). We analyzed this data for expression of *ALDOB* and found that the transcript levels of *ALDOB* did not change significantly between normal and obese individuals. However, there was a dramatic down regulation of ALDOB mRNA in NAFLD (2.08 fold, p<0.001) and NASH (2.9 fold, p<0.001) patients (Fig. 4D). This gives credence to the idea that ALODB down regulation might be an important and novel pathophysiological mechanism for fatty liver disease.

## Discussion

NAFLD is known to be associated with obesity, however there is mounting evidence for NAFLD in lean individuals, especially in Asian populations (5). Lean NAFLD appears to be more common in older, male and Asian individuals compared to other groups (43). A comparative study of lean healthy individuals from various ethnicities found that Asian-Indian men had higher hepatic triglyceride levels and a higher basal serum IL6 (6). As discussed in the Introduction, many autoimmune diseases characterized by elevated circulating IL6 such as SLE, Crohn’s disease, IBD and RA also appear to have high propensity for fatty liver (14-21). Various other studies also show elevated inflammation and IL6 expression in NAFLD although this is considered to be more a consequence than cause (10, 44). Taken together, these studies hint at a tantalizing relationship between inflammation and fatty liver. In this study for the first time, we report a causative role for IL6 in the development of non-alcoholic fatty liver disease. We demonstrate that chronic systemic exposure to IL6 in zebrafish induces hepatic steatosis selectively in the male.

A number of studies on animal models for HFD-induced NAFLD and NAFLD patient samples have revealed the molecular changes that lead to fat accumulation in the liver. In these conventional models of NAFLD there is evidence for up regulation of lipid biosynthetic pathway genes such as *srebp1, pparab, pparg, fads2, and cd36* (45, 46). However, our inflammation model does not show these expected changes in the liver suggesting a different pathophysiology for fatty liver disease induced by inflammation.

One of the consistent changes we observed in our RNA sequencing studies was down regulation of the glycolysis/gluconeogenesis pathway specifically in the male IL6-OE zebrafish liver. Of the glycolysis genes, the enzyme Aldolase b was consistently down regulated in both our transcriptomics as well as proteomics data. Treatment of HepG2 cells with recombinant IL6 also led to the suppression of ALDOB expression. Our RNA sequencing experiments revealed a down regulation of PPAR signaling components, similar to that seen in NAFLD patients (33). Previous studies have predicted PPAR response elements on the promoter of *ALDOB* (47) suggesting transcriptional regulation of *aldolase b* gene by PPAR signaling pathway in the IL6-OE zebrafish..This would need further investigation.

Aldolase b converts fructose-1,6-bisphosphate into dihydroxy acetone phosphate and glyceraldehyde-3-phosphate in the glycolytic pathway. Mice null for Aldob are intolerant to fructose in diet (48). The rare inborn error of metabolism known as hereditary fructose intolerance (HFI) in humans is caused by mutations in the *ALDOB* gene (49). HFI is also characterized by the accumulation of mono-glyco:di-glyco. Consistent with these studies, the male IL6-OE zebrafish liver showed an accumulation of hexose monophosphates. These observations are of special interest because a very recent study found that patients with ALDOB deficiency have elevated intrahepatic triglyceride content. Our metabolomics analysis offers an insight into a possible mechanism. Our data suggests that deregulation of glycolysis/gluconeogenesis can lead to an accumulation of DHAP/glyceraldehyde-3-phosphate, which is then converted into triglycerides via sn-glycerol-3-phosphate in the IL6-OE liver.

The ALDOB deficient HFI patients with fatty liver have normal BMI categorizing these patients as lean fatty liver. NAFLD in lean patients, a phenomenon particularly noticeable in Indian males, remains intriguing. We hypothesize that chronic systemic inflammation, particularly elevated levels of IL6, can trigger intrahepatic triglyceride and NAFLD like disease. We further speculate that IL6 induces NAFLD-like phenotype in lean individuals through metabolic deregulation of glycolysis/gluconeogenesis and channeling of metabolites into triglyceride synthesis. This proposed molecular mechanism of triglyceride accumulation in deregulated sugar metabolism would be an interesting avenue for pursuit.

## List of Abbreviations

NAFLD: Non-Alcoholic Fatty Liver Disease
IL6: Interleukin 6
OE: Overexpression
HFI: High Fructose Intolerance
NASH: Non-Alcoholic Steatohepatitis
SLE: Systemic Lupus Erythematosus
IBD: Inflammatory Bowel Disease
RA: Rheumatoid Arthritis
HFD: High Fat diet
qRT-PCR: Quantitative Real-Time Polymerase Chain Reaction
PPAR: Peroxisome Proliferator-Activated Receptor
DHAP: Dihydroxyacetone Phosphate

## Acknowledgments

We appreciate the help of Rohit Yadav, Helly Shah and Shashi Ranjan with zebrafish experiments. We acknowledge Amit Chandra for helping with western blot standardization and Farina Sultan for help with microscopy. We acknowledge Manish for help with confocal experiments and the TEM central facility for ultra-structural studies. We acknowledge Monika Verma and Shweta Verma for final proofreading.

## Notes

**Funding Information:** This work was supported by Council of Scientific and Industrial Research (CSIR), New Delhi [BSC0124, MLP1801]. M.K.S was supported by Council of Scientific and Industrial Research (CSIR) research fellowship. The funders had no role in study design, data collection and analysis, decision to publish, or preparation of the article.

### Competing Interest Statement

The authors have declared no competing interest.

## References

1. Younossi ZM. Non-alcoholic fatty liver disease - A global public health perspective. J Hepatol 2019;70:531–544.

2. Mitra S, De A, Chowdhury A. Epidemiology of non-alcoholic and alcoholic fatty liver diseases. Transl Gastroenterol Hepatol 2020;5:16.

3. D’Avola D, Labgaa I, Villanueva A. Natural history of nonalcoholic steatohepatitis/nonalcoholic fatty liver disease-hepatocellular carcinoma: Magnitude of the problem from a hepatology clinic perspective. Clin Liver Dis (Hoboken) 2016;8:100–104.

4. Ahmed MH, Husain NE, Almobarak AO. Nonalcoholic Fatty liver disease and risk of diabetes and cardiovascular disease: what is important for primary care physicians? J Family Med Prim Care 2015;4:45–52.

5. Albhaisi S, Chowdhury A, Sanyal AJ. Non-alcoholic fatty liver disease in lean individuals. JHEP Rep 2019;1:329–341.

6. Petersen KF, Dufour S, Feng J, Befroy D, Dziura J, Dalla Man C, Cobelli C, et al. Increased prevalence of insulin resistance and nonalcoholic fatty liver disease in Asian-Indian men. Proc Natl Acad Sci U S A 2006;103:18273–18277.

7. Joshi-Barve S, Barve SS, Amancherla K, Gobejishvili L, Hill D, Cave M, Hote P, et al. Palmitic acid induces production of proinflammatory cytokine interleukin-8 from hepatocytes. Hepatology 2007;46:823–830.

8. Cui W, Chen SL, Hu KQ. Quantification and mechanisms of oleic acid-induced steatosis in HepG2 cells. Am J Transl Res 2010;2:95–104.

9. Lieber CS, Leo MA, Mak KM, Xu Y, Cao Q, Ren C, Ponomarenko A, et al. Model of nonalcoholic steatohepatitis. Am J Clin Nutr 2004;79:502–509.

10. Wieckowska A, Papouchado BG, Li Z, Lopez R, Zein NN, Feldstein AE. Increased hepatic and circulating interleukin-6 levels in human nonalcoholic steatohepatitis. Am J Gastroenterol 2008;103:1372–1379.

11. Paredes-Turrubiarte G, Gonzalez-Chavez A, Perez-Tamayo R, Salazar-Vazquez BY, Hernandez VS, Garibay-Nieto N, Fragoso JM, et al. Severity of non-alcoholic fatty liver disease is associated with high systemic levels of tumor necrosis factor alpha and low serum interleukin 10 in morbidly obese patients. Clin Exp Med 2016;16:193–202.

12. Palomera LF, Gomez-Arauz AY, Villanueva-Ortega E, Melendez-Mier G, Islas-Andrade SA, Escobedo G. Serum levels of interleukin-1 beta associate better with severity of simple steatosis than liver function tests in morbidly obese patients. J Res Med Sci 2018;23:93.

13. Yuan J, Chen C, Cui J, Lu J, Yan C, Wei X, Zhao X, et al. Fatty Liver Disease Caused by High-Alcohol-Producing Klebsiella pneumoniae. Cell Metab 2019;30:675–688 e677.

14. Bessissow T, Le NH, Rollet K, Afif W, Bitton A, Sebastiani G. Incidence and Predictors of Nonalcoholic Fatty Liver Disease by Serum Biomarkers in Patients with Inflammatory Bowel Disease. Inflamm Bowel Dis 2016;22:1937–1944.

15. Ripley BJ, Goncalves B, Isenberg DA, Latchman DS, Rahman A. Raised levels of interleukin 6 in systemic lupus erythematosus correlate with anaemia. Ann Rheum Dis 2005;64:849–853.

16. Reinisch W, Gasche C, Tillinger W, Wyatt J, Lichtenberger C, Willheim M, Dejaco C, et al. Clinical relevance of serum interleukin-6 in Crohn’s disease: single point measurements, therapy monitoring, and prediction of clinical relapse. Am J Gastroenterol 1999;94:2156–2164.

17. Madhok R, Crilly A, Watson J, Capell HA. Serum interleukin 6 levels in rheumatoid arthritis: correlations with clinical and laboratory indices of disease activity. Ann Rheum Dis 1993;52:232–234.

18. Matsumoto T, Kobayashi S, Shimizu H, Nakajima M, Watanabe S, Kitami N, Sato N, et al. The liver in collagen diseases: pathologic study of 160 cases with particular reference to hepatic arteritis, primary biliary cirrhosis, autoimmune hepatitis and nodular regenerative hyperplasia of the liver. Liver 2000;20:366–373.

19. Bessone F, Poles N, Roma MG. Challenge of liver disease in systemic lupus erythematosus: Clues for diagnosis and hints for pathogenesis. World J Hepatol 2014;6:394–409.

20. Sartini A, Gitto S, Bianchini M, Verga MC, Di Girolamo M, Bertani A, Del Buono M, et al. Non-alcoholic fatty liver disease phenotypes in patients with inflammatory bowel disease. Cell Death Dis 2018;9:87.

21. Abraham S, Begum S, Isenberg D. Hepatic manifestations of autoimmune rheumatic diseases. Ann Rheum Dis 2004;63:123–129.

22. Ma KL, Ruan XZ, Powis SH, Chen Y, Moorhead JF, Varghese Z. Inflammatory stress exacerbates lipid accumulation in hepatic cells and fatty livers of apolipoprotein E knockout mice. Hepatology 2008;48:770–781.

23. Endo M, Masaki T, Seike M, Yoshimatsu H. TNF-alpha induces hepatic steatosis in mice by enhancing gene expression of sterol regulatory element binding protein-1c (SREBP-1c). Exp Biol Med (Maywood) 2007;232:614–621.

24. Asad Z, Pandey A, Babu A, Sun Y, Shevade K, Kapoor S, Ullah I, et al. Rescue of neural crest-derived phenotypes in a zebrafish CHARGE model by Sox10 downregulation. Hum Mol Genet 2016;25:3539–3554.

25. Steinbicker AU, Sachidanandan C, Vonner AJ, Yusuf RZ, Deng DY, Lai CS, Rauwerdink KM, et al. Inhibition of bone morphogenetic protein signaling attenuates anemia associated with inflammation. Blood 2011;117:4915–4923.

26. Pandey A, Ekka MK, Ranjan S, Maiti S, Sachidanandan C. Teratogenic, cardiotoxic and hepatotoxic properties of related ionic liquids reveal the biological importance of anionic components. RSC Advances 2017;7:22927–22935.

27. Pfaffl MW. A new mathematical model for relative quantification in real-time RT-PCR. Nucleic Acids Res 2001;29:e45.

28. Basu S, Jalodia K, Ranjan S, Yeh JJ, Peterson RT, Sachidanandan C. Small Molecule Inhibitors of NFkB Reverse Iron Overload and Hepcidin Deregulation in a Zebrafish Model for Hereditary Hemochromatosis Type 3. ACS Chem Biol 2018;13:2143–2152.

29. Schneider CA, Rasband WS, Eliceiri KW. NIH Image to ImageJ: 25 years of image analysis. Nat Methods 2012;9:671–675.

30. Isobe A, Takeda T, Sakata M, Yamamoto T, Minekawa R, Hayashi M, Auernhammer CJ, et al. STAT3-mediated constitutive expression of SOCS3 in an undifferentiated rat trophoblast-like cell line. Placenta 2006;27:912–918.

31. Lambert JE, Ramos-Roman MA, Browning JD, Parks EJ. Increased de novo lipogenesis is a distinct characteristic of individuals with nonalcoholic fatty liver disease. Gastroenterology 2014;146:726–735.

32. Dorn C, Riener MO, Kirovski G, Saugspier M, Steib K, Weiss TS, Gabele E, et al. Expression of fatty acid synthase in nonalcoholic fatty liver disease. Int J Clin Exp Pathol 2010;3:505–514.

33. Francque S, Verrijken A, Caron S, Prawitt J, Paumelle R, Derudas B, Lefebvre P, et al. PPARalpha gene expression correlates with severity and histological treatment response in patients with non-alcoholic steatohepatitis. J Hepatol 2015;63:164–173.

34. Wang W, Zhang X, Qin J, Wei P, Jia Y, Wang J, Ru S. Long-term bisphenol S exposure induces fat accumulation in liver of adult male zebrafish (Danio rerio) and slows yolk lipid consumption in F1 offspring. Chemosphere 2019;221:500–510.

35. Diao L, Auger C, Konoeda H, Sadri AR, Amini-Nik S, Jeschke MG. Hepatic steatosis associated with decreased beta-oxidation and mitochondrial function contributes to cell damage in obese mice after thermal injury. Cell Death Dis 2018;9:530.

36. Lebeaupin C, Vallee D, Hazari Y, Hetz C, Chevet E, Bailly-Maitre B. Endoplasmic reticulum stress signalling and the pathogenesis of non-alcoholic fatty liver disease. J Hepatol 2018;69:927–947.

37. Zheng X, Dai W, Chen X, Wang K, Zhang W, Liu L, Hou J. Caffeine reduces hepatic lipid accumulation through regulation of lipogenesis and ER stress in zebrafish larvae. J Biomed Sci 2015;22:105.

38. Langmead B, Salzberg SL. Fast gapped-read alignment with Bowtie 2. Nat Methods 2012;9:357–359.

39. Trapnell C, Williams BA, Pertea G, Mortazavi A, Kwan G, van Baren MJ, Salzberg SL, et al. Transcript assembly and quantification by RNA-Seq reveals unannotated transcripts and isoform switching during cell differentiation. Nat Biotechnol 2010;28:511–515.

40. Huang da W, Sherman BT, Lempicki RA. Systematic and integrative analysis of large gene lists using DAVID bioinformatics resources. Nat Protoc 2009;4:44–57.

41. Simons N, Debray FG, Schaper NC, Kooi ME, Feskens EJM, Hollak CEM, Lindeboom L, et al. Patients With Aldolase B Deficiency Are Characterized by Increased Intrahepatic Triglyceride Content. J Clin Endocrinol Metab 2019;104:5056–5064.

42. Suppli MP, Rigbolt KTG, Veidal SS, Heeboll S, Eriksen PL, Demant M, Bagger JI, et al. Hepatic transcriptome signatures in patients with varying degrees of nonalcoholic fatty liver disease compared with healthy normal-weight individuals. Am J Physiol Gastrointest Liver Physiol 2019;316:G462–G472.

43. Zou B, Yeo YH, Nguyen VH, Cheung R, Ingelsson E, Nguyen MH. Prevalence, characteristics and mortality outcomes of obese, nonobese and lean NAFLD in the United States, 1999-2016. J Intern Med 2020.

44. Haukeland JW, Damas JK, Konopski Z, Loberg EM, Haaland T, Goverud I, Torjesen PA, et al. Systemic inflammation in nonalcoholic fatty liver disease is characterized by elevated levels of CCL2. J Hepatol 2006;44:1167–1174.

45. Dai W, Wang K, Zheng X, Chen X, Zhang W, Zhang Y, Hou J, et al. High fat plus high cholesterol diet lead to hepatic steatosis in zebrafish larvae: a novel model for screening anti-hepatic steatosis drugs. Nutr Metab (Lond) 2015;12:42.

46. Park KH, Ye ZW, Zhang J, Kim SH. Palmitic Acid-Enriched Diet Induces Hepatic Steatosis and Injury in Adult Zebrafish. Zebrafish 2019;16:497–504.

47. Lemay DG, Hwang DH. Genome-wide identification of peroxisome proliferator response elements using integrated computational genomics. J Lipid Res 2006;47:1583–1587.

48. Oppelt SA, Sennott EM, Tolan DR. Aldolase-B knockout in mice phenocopies hereditary fructose intolerance in humans. Mol Genet Metab 2015;114:445–450.

49. Santer R, Rischewski J, von Weihe M, Niederhaus M, Schneppenheim S, Baerlocher K, Kohlschutter A, et al. The spectrum of aldolase B (ALDOB) mutations and the prevalence of hereditary fructose intolerance in Central Europe. Hum Mutat 2005;25:594.

