## Supplementary figures and images for "Chronic exposure to IL6 leads to deregulation of glycolysis and fat accumulation in the zebrafish liver"

### Supplementary Figure 1

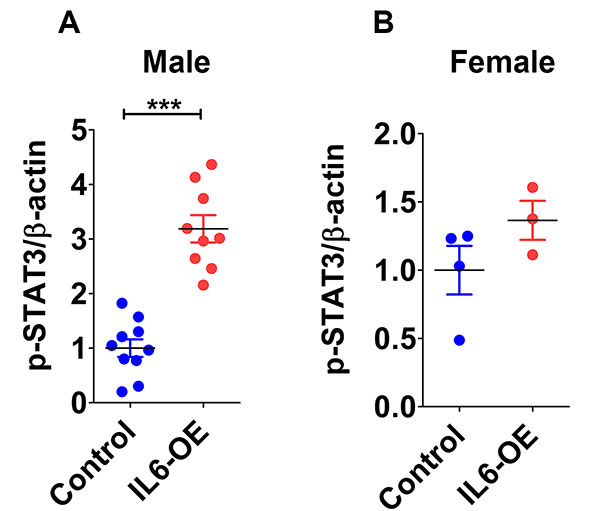

### Supplementary Figure 2

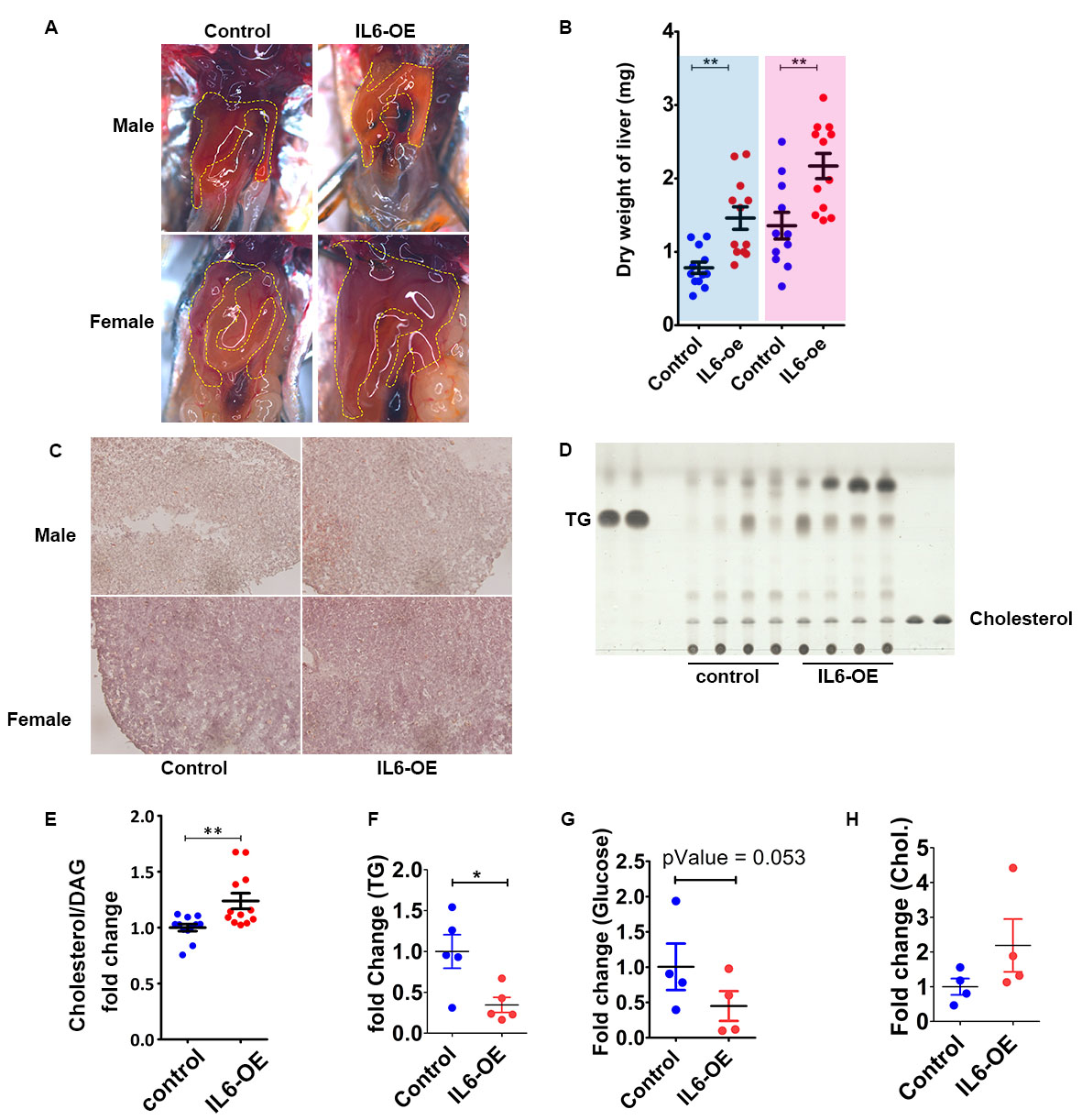

### Supplementary Figure 3

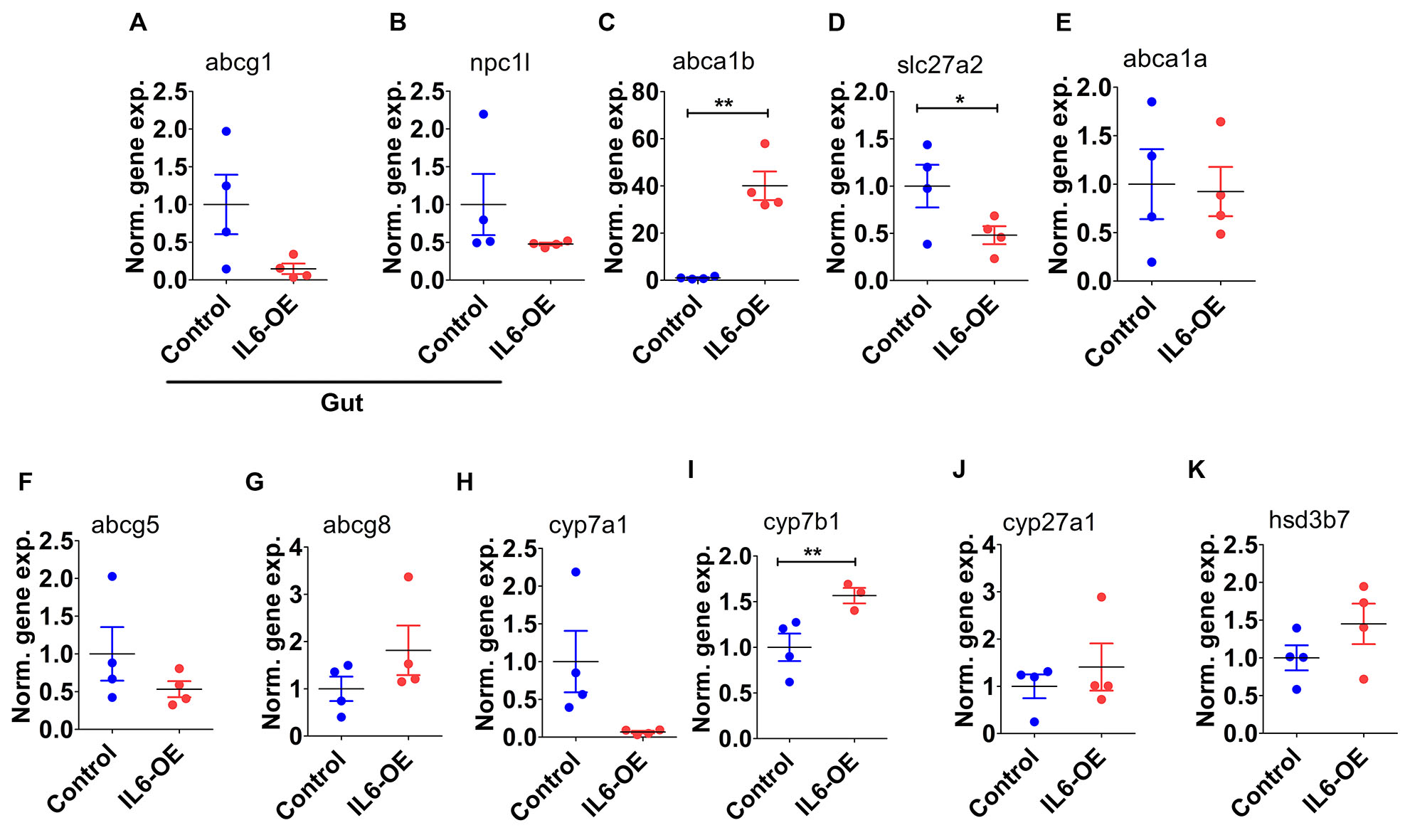

### Supplementary Figure 4

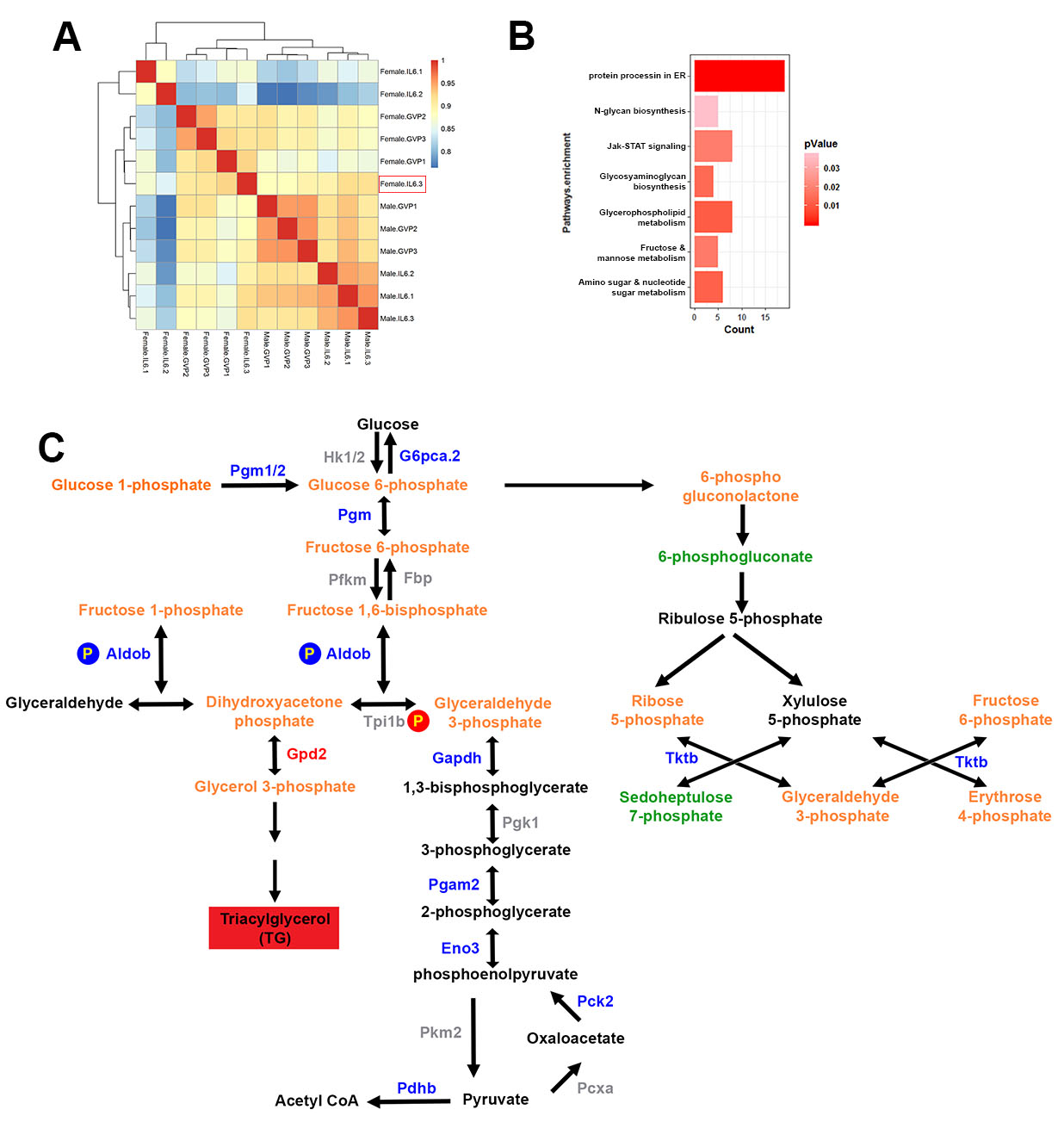

### Supplementary Figure 5

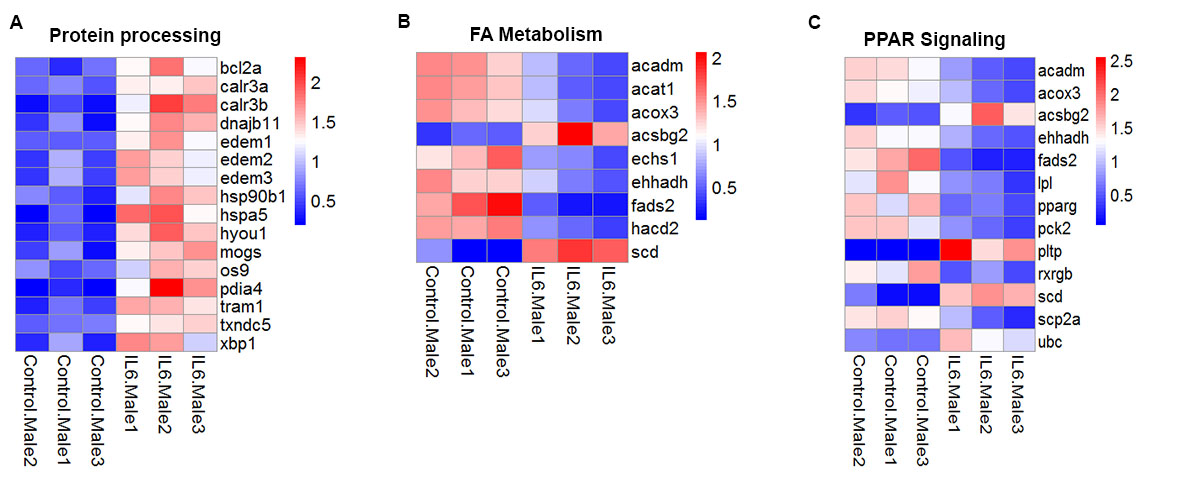

### Supplementary Figure 6

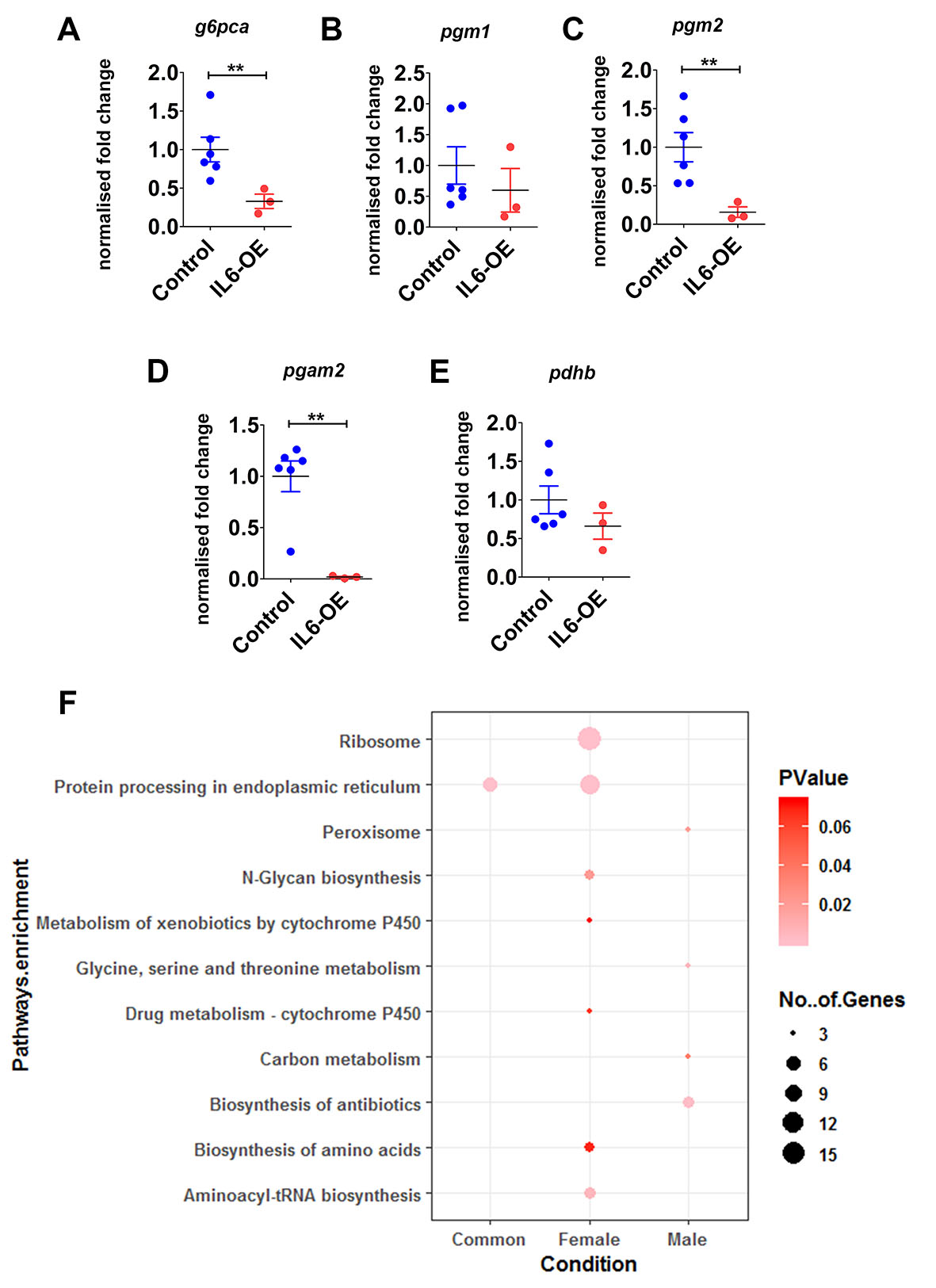

### Supplementary Figure 7

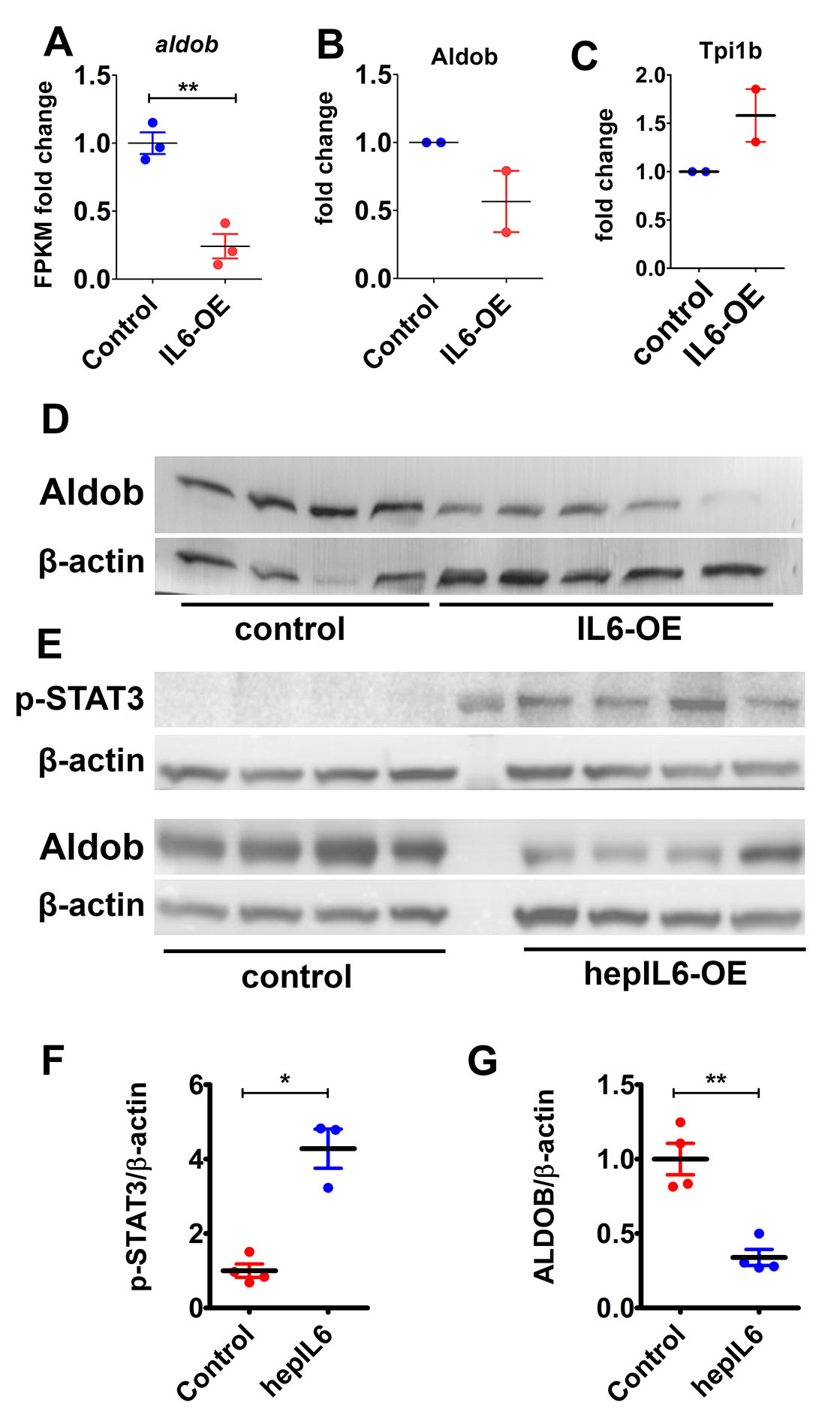

### Supplementary Figure 8

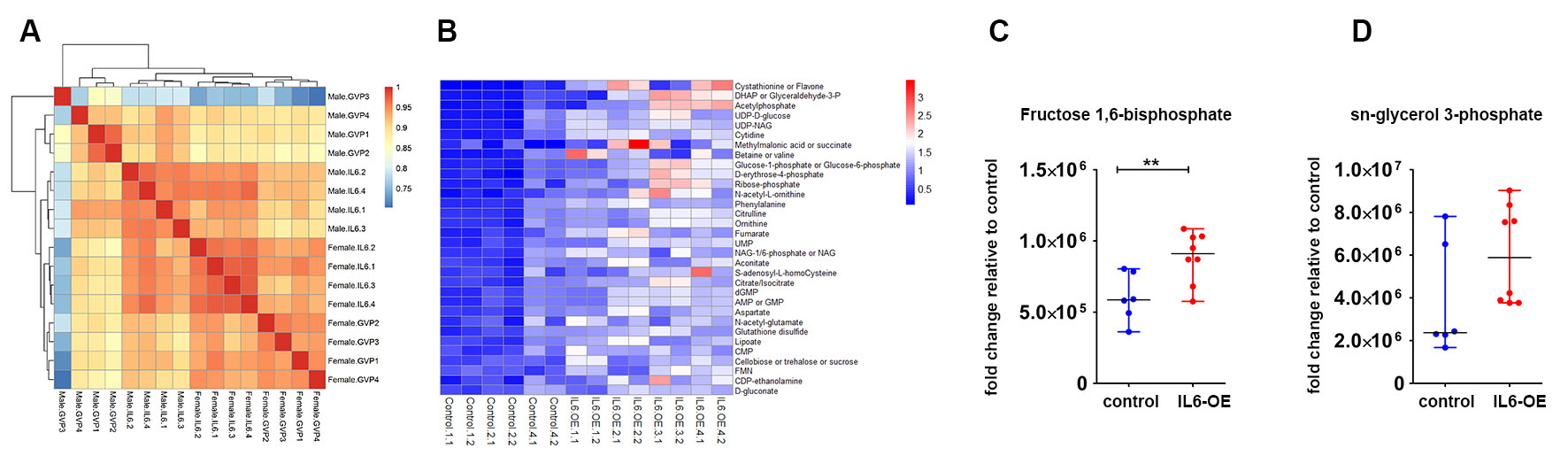
