## Supplementary Methods and Legends for "Chronic exposure to IL6 leads to deregulation of glycolysis and fat accumulation in the zebrafish liver"

**SUPPLEMENTARY INFORMATION**

**Supplementary Methods**

**WESTERN BLOT ANALYSIS**

Adult zebrafish liver and heart tissue were homogenized in NP40 lysis buffer (Thermo Fisher, FNN0021). The lysates were centrifuged (14000 rpm for 20 min at 4 °C), and supernatants were collected. The protein concentration was estimated using BCA (Pierce 23225) method. Total protein (25 μg) was resolved on a 12% gel using SDS-PAGE method. The proteins were transferred on a PVDF membrane (Merck Millipore, ISEQ00010), and 5% BSA in Tris-buffered saline (TBS) containing 0.1% (v/v) Tween-20 (TBST) was used for blocking the membranes for 1h. Incubation with 1:1000 diluted primary antibodies against h-IL6 (ab6672), Aldolase B (GTX124306), Stat3 (4904s), Stat3-phospho (9131s) was done overnight at 4 °C. Following, the blots were washed thrice with TBST and incubated with 1:10000 diluted HRP conjugate secondary anti rabbit IgG (Cell Signaling, 7074S) antibody for 1.5 h. The signal was detected using EMD Millipore Immobilon Western Chemiluminescent HRP Substrate (WBKLS0500) on a G:BOX detection platform (Syngene).

**TRANSCRIPTOMICS**

RNA was isolated from adult liver (from 3 animals in each group) of control males and females and IL6-OE males and females. RNA-seq libraries were prepared from 1µg of total RNA using standard protocol. Paired end method for RNA sequencing was performed on HiSeq2500 sequencing platform from Illumina, USA. We obtained an average depth of 52 million reads. The quality of the reads were checked using FastQC and the reads were trimmed for adapters and then sorted using Trimmomatic using a quality score of 20. Reads were aligned to the reference genome using TopHat (Genome aligner Bowtie-2-2.1.0). The reads were annotated using cufflinks which were quantified and normalised with cuffnorm. We mapped the reads to the zebrafish reference genome (Zv11) and achieved >60-89% alignment using Bowtie tool (1). The transcripts were annotated using the reference transcript of ENSEMBL v87 and FPKM of each gene from all the samples were calculated using cuffnorm (2). The FPKM value of genes from IL6-OE male and female livers were compared to their respective controls. We removed all genes with an FPKM=0 in any of the samples to simplify the analysis. The raw data containing reads were uploaded to NCBI SRA database with the Bioproject accession ID: PRJNA638724

**PROTEOMICS**

Liver samples of WT males (n = 2), WT females (n = 2), IL6-OE males (n = 2) and IL6-OE females (n = 2) were homogenized in a lysis buffer containing 7 M urea, 2 M thiourea, 4% (w/v) CHAPS and EDTA-free 1× protease inhibitor cocktail. The lysates were centrifuged at 13,000 rpm for 15 min at 4 °C, and the supernatant were collected and stored at −80 °C. Total protein concentration was estimated using Bradford reagent. Total protein (50 μg) from each sample were digested overnight with trypsin at 37 °C. The protein were labelled using iTRAQ reagent as per the manufacturer’s protocol (AB Sciex). In brief, iTRAQ labeling tags (113, 114, 115, 116, 117, 118, 119, and 121) were reconstituted in 50 µL absolute isopropanol and were added to the 8 samples respectively. The samples tagged with iTRAQ were pooled in a single vial and dried using a vacuum concentrator. The dried sample was then fractionated using the strong cation exchange fractionation an ICAT Cartridge (AB Sciex, Foster City, CA, USA) and peptides were eluted with the sequential concentration of 35 mM, 75 mM, 100 mM, 125 mM, 150 mM, 250 mM, 350 mM, and 500 mM ammonium formate buffer, pH 3. Lastly, for subsequent LC-MS/MS analysis, samples were reconstituted in 95%water 5% acetonitrile, 0.1% formic acid and were analyzed on a 5600 triple time-of-flight (TOF) (AB Sciex) coupled with Eksigent nano-LC (3). The fractions were separated on reverse phase C18 analytical column (ChromXP 75 μm × 15 cm, 3 μm 120 Å) and with trap column (ChromXP 350 μm × 0.5 mm, 3 μm 120 Å). The triple TOF 5600 (AB Sciex) was operated in data-dependent acquisition mode and full MS spectra were acquired in positive ion mode in m/z 350-1200 Da, whereas the MS/MS product was performed in the mass range of 100-2000 Da. Raw data files were searched against the Zebrafish reference proteomes (UniProt Proteome ID: UP000000437) using Protein Pilot (version 4.0). The proteins were considered differentially expressed if the ratio was > 1.2-fold or < 0.8-fold in both the replicates of male and female and was identified with a minimum of 2 peptides.

**LIVER METABOLOMICS**

Before sample extraction, the cryofixed livers were weighed and solvent volume used for extraction was adjusted to normalize for weight difference between samples such that every 1mg of tissue was extracted with 200uL of solvent. Extraction was carried out as previously reported (4). Briefly, the liver samples were resuspended in ice cold 90% methanol water solution (containing 1ng/ml PIPES as internal standard) and incubated on ice for 10 mins. The samples were then sonicated for 15 minutes in an iced water bath and centrifuged for 5 minutes at 14,000 rpm the the supernatant was collected. This procedure was repeated twice more for efficient extraction of metabolites and extracts of same sample were pooled and stored in -80°C until further LC-MS analysis. 10uL of the each sample extract was run on a Thermo Q-Exactive Orbitrap mass spectrometer coupled to an Accela U-HPLC (Thermo Fisher Scientific) fitted with HTC PAL autosampler (CTC Analytics AG, Switzerland). Thermo Accucore RP C18 column with a bed volume of 150 mm x 2.1 mm and 2.6 μ particle size was used for separation of metabolites. The column flow rate was maintained at 400uL/min and solvent system composed of acetonitrile buffered with 0.1% formic acid (buffer A) and water buffered with 0.1% formic acid (buffer B) was used in a 16 min gradient run with the following parameters: Linear increase in buffer A till it reaches 95%. Hold at 95% buffer A for 2 mins and then ramp down till it reaches 1% and hold for 2 mins. Metabolite ions between 70-1000 mass-charge ratio (m/z value) were monitored and data collected in negative ionization mode.

**LC-MS data processing and analysis:**

The raw data files were converted into .mzXML file format using the Proteowizard tool (<http://proteowizard.sourceforge.net/>) and further data analysis and visualization were carried out using the El-MAVEN software (5) to identify the various metabolites of interest within 10 ppm error on mass. The list containing the metabolites of interest and their fold change values is available in the Supporting Table S4. Male control (male GVP3) was found to be an outlier and did not cluster with the other samples of its group and was excluded from data analysis (Supplementary Fig. 8A).

**References:**

1.	Langmead B, Salzberg SL. Fast gapped-read alignment with Bowtie 2. Nat Methods 2012;9:357-359.

2.	Trapnell C, Williams BA, Pertea G, Mortazavi A, Kwan G, van Baren MJ, Salzberg SL, et al. Transcript assembly and quantification by RNA-Seq reveals unannotated transcripts and isoform switching during cell differentiation. Nat Biotechnol 2010;28:511-515.

3.	Varshney S BN, Basak T, Sengupta S. Identification of differentially expressed proteins in vitamin B 12. J Pract Cardiovasc Sci 2015;1:45-53.

4.	Shukla A, Olszewski KL, Llinas M, Rommereim LM, Fox BA, Bzik DJ, Xia D, et al. Glycolysis is important for optimal asexual growth and formation of mature tissue cysts by Toxoplasma gondii. Int J Parasitol 2018;48:955-968.

5.	Agrawal S, Kumar S, Sehgal R, George S, Gupta R, Poddar S, Jha A, et al. El-MAVEN: A Fast, Robust, and User-Friendly Mass Spectrometry Data Processing Engine for Metabolomics. Methods Mol Biol 2019;1978:301-321.

**Supplementary Figure Legends:**

**Supplementary Figure 1. Induction of hepatic STAT-3 phosphorylation IL6-OE zebrafish** (A) Western blot (Figure 1E) quantification showed significant induction of Stat-3 phosphorylation in IL6-OE liver as compared to control liver of male zebrafish. (B) Western blot (Figure 1E) quantification showed increase of Stat-3 phosphorylation in IL6-OE as compared to control female zebrafish. Data presented as mean ± SE. *p* value is calculated using unpaired student t-test, *** p< 0.001. Each point in graph represents a single animal.

**Supplementary Figure 2. IL6 overexpression causes morphological changes in male zebrafish.** (A) Morphological changes were observed in IL6-OE liver in male zebrafish as compared to control male, control female and IL6-OE female. The boundaries of the liver is marked by yellow dotted outline. Images were taken at magnification of 1.25X. (B) Dry weight of liver was significantly increased in IL6-OE male and female as compared to their respective controls. Data in blue box is for male and pink box is for female. (C) Neutral lipid stained using Oil red O shows lipid accumulation in male IL6-OE. No significant lipid accumulation was evident in the other samples. (D) Thin layer chromatography with two standards triglyceride (TG) and cholesterol. The solvents used were suitable for TG for separation. The band co-migrating with the cholesterol standard could be choesterol and/or diacylglycerol (DAG). (E) Quantification of DAG/cholesterol shows significant accumulation in IL6-OE male liver. The standards for triglyceride and cholesterol are indicated. (F-H) Serum biochemical analysis for triglyceride, glucose and cholesterolshowed significant reduction of triglyceride in the male IL6-OE zebrafish liver. Data presented as mean ± SE. *p* value is calculated using unpaired student t-test, * p< 0.05 and ** p< 0.01. Each point in graph represents a single animal.

**Supplementary Figure 3. Expression of genes involved in lipid transport and bile acid synthesis by qRT-PCR.** (A-B) Gene expression fold change normalized to beta-actin. No significant change in npc1l1 and abcg1 in gut tissue of IL6-OE animals. (C-E) Cholesterol transport gene *abca1b* was induced and *slc27a2* was down regulated in male IL6-OE. Fold change normalized to *rpl13*α. (F-K) Expression of genes involved in bile acid synthesis and secretion. *cyp7b1* shows significant up regulation in male IL6-OE liver. No significant changes were found in *abcg5, abcg8, cyp7a1, cyp27a1* and *hsd3b7*. Fold change normalized to *rpl13*α. Data presented as mean ± SE. *p* value is calculated using unpaired student t-test, * p< 0.05 and ** p< 0.01. Each point in graph represents a single animal.

**Supplementary Figure 4. RNA Sequencing of IL6-OE adult liver reveals enrichment of pathways.** (A) Spearman correlation revealed that Female.IL6.3 sample (red box) clustered separately from other female IL6-OE samples. This sample was removed from further analysis. (B) Genes up regulated in male IL6-OE show KEGG pathway enrichment for protein processing in ER and JAK-STAT signaling. Count represents the number of genes enriched in the particular pathways. (C) Glycolysis/gluconeogenesis pathway with the enzymes involved in each step. The enzymes in blue are down regulated in the IL6-OE male liver transcriptomics. Enzymes in grey are not part of the 791 DEGs. The red circles represent up regulation and blue circles represent down regulation in the proteomics experiment.

**Supplementary Figure 5. KEGG pathway enrichment analysis of DEGs in IL6-OE male.** (A) A number of genes involved in protein processing in ER show up regulation in IL6-OE male. (B-C) Genes involved in fatty acid metabolism and PPAR signaling show down regulation in IL6-OE male.

**Supplementary Figure 6. Glycolysis and carbon metabolism down regulated in male IL6-OE liver. (A-E)** qRT-PCR validation of down regulation of glycolysis genes, *g6pca*, *pgm1*, *pgm2*, *pgam2* and *pdhb* in male IL6-OE liver. Fold change normalized to *rpl13*α. The data is presented as mean ± SE. *p* value is calculated using unpaired student t-test, ** p< 0.01. Each point in graph represents single animal. (F) Proteomics profiling shows enrichment of carbon metabolic pathway in male IL6-OE liver and ribosomal genes in female IL6-OE.

**Supplementary Figure 7. Overexpression of humanIL6 in the liver leads to down regulation of hepatic *aldob* in IL6-OE male.** (A) Fold change of *aldob* mRNA FPKM showed significant down regulation in IL6-OE male. (B-C) Fold change of Aldob and Tpi1b protein quantified using protein sequencing shows down regulation and up regulation in IL6-OE male respectively. (D) Representative western blot analysis for Aldob in control and IL6-OE male samples. (E) Western blot analysis of liver protein from hepatic overexpression of humanIL6 (hepIL6-OE) induces STAT-3 phosphorylation and down regulation of Aldob protein levels in IL6-OE male. (F-G) Quantification of western blot signal using Image J software shows statistically significant up regulation of p-STAT3 and down regulation of Aldob in IL6-OE male liver. The data is presented as mean ± SE. *p* value is calculated using unpaired student t-test, * p< 0.05 and ** p< 0.01. Each point in graphs represents single animal.

**Supplementary Figure 8. Differential regulation of metabolites in male IL6-OE liver.** (A) Spearman correlation analysis of metabolic profiling experiment showed one sample (Male.GVP3) clustering separately from other samples of the group. This sample was removed from further analysis. (B) Differential accumulation of metabolites in IL6-OE male liver. Each biological sample is run in two technical replicates. (C-D) Accumulation of fructose-1,6-bisphosphate and sn-glycerol-3-phophate in IL6-OE male liver. The data is presented as median ± range. *p* value is calculated using unpaired student t-test, ** p< 0.01. Points represent both biological as well as technical replicates.
